# SynapseJ: an automated, synapse identification macro for ImageJ

**DOI:** 10.1101/2021.06.24.449851

**Authors:** Juan Felipe Moreno Manrique, Parker R. Voit, Kathryn E. Windsor, Aamuktha R. Karla, Sierra R. Rodriguez, Gerard M. J. Beaudoin

## Abstract

While electron microscopy represents the gold standard for detection of synapses, a number of limitations prevent its broad applicability. A key method for detecting synapses is immunostaining for markers of pre- and post-synaptic proteins, which can infer a synapse based upon the apposition of the two markers. While immunostaining and imaging techniques have improved to allow for identification of synapses in tissue, analysis and identification of these appositions are not facile, and there has been a lack of tools to accurately identify these appositions. Here, we delineate a macro that uses open-source and freely available ImageJ or FIJI for analysis of multichannel, z-stack confocal images. With use of a high magnification with a high NA objective, we outline two methods to identify puncta in either sparsely or densely labeled images. Puncta from each channel are used to eliminate non-apposed puncta and are subsequently linked with their cognate from the other channel. These methods are applied to analysis of a presynaptic marker, bassoon, with two different postsynaptic markers, gephyrin and N-methyl-d-aspartate (NMDA) receptor subunit 1 (NR1). Using gephyrin as an inhibitory, postsynaptic scaffolding protein, we identify inhibitory synapses in basolateral amygdala, central amygdala, arcuate and the ventromedial hypothalamus. Systematic variation of the settings identify the parameters most critical for this analysis. Identification of specifically overlapping puncta allows for correlation of morphometry data between each channel. Finally, we extend the analysis to only examine puncta overlapping with a cytoplasmic marker of specific cell types, a distinct advantage beyond electron microscopy. Bassoon puncta are restricted to virally transduced, pedunculopontine tegmental neuron (PPN) axons expressing yellow fluorescent protein. NR1 puncta are restricted to tyrosine hydroxylase labeled dopaminergic neurons of the substantia nigra pars compacta (SNc). The macro identifies bassoon-NR1 overlap throughout the image, or those only restricted to the PPN-SNc connections. Thus, we have extended the available analysis tools that can be used to study synapses in situ. Our analysis code is freely available and open-source allowing for further innovation.

## 1 Introduction

Quantification of digitally acquired images is a field standard for any microscopic data (For example: Beaudoin et al., 2012;Taylor et al., 2020;Pourhoseini et al., 2021). Many kinds of information lend themselves to simple means of quantification, including dendritic complexity being analyzed by Sholl analysis, protein expression level by intensity analysis, or colocalization of two proteins by correlation analysis (Beaudoin et al., 2012;Nelson et al., 2016;Boulan et al., 2020;Ligon et al., 2020). However, identification of chemical synapses, henceforth referenced to simply as a synapse, from imaging data represents a unique challenge. The standard has been to rely on electron microscopy for accurate identification of synapses, and this analysis was done visually by a user, though this is increasingly being done by computers (Merchan-Perez et al., 2009;Beaudoin et al., 2012;Kornfeld and Denk, 2018;Nosov et al., 2020;Pourhoseini et al., 2021). The unique challenge of synaptic identification is that a synapse is defined as the zone of interaction between two neurons, where one neuron releases neurotransmitter and the other neuron receives the neurotransmitter. To identify these two components using fluorescent microscopy requires the use of antibodies that selectively identify a protein from each compartment.

The identification of a protein that selectively and specifically targets a pre-synaptic or post-synaptic structure requires careful consideration. Synapses were originally identified by proteins found on synaptic vesicles, a necessary component of a synapse (Bixby and Reichardt, 1985). However, synaptic vesicles are known to be in parts of the axon away from the release site (Staras and Branco, 2010). Additionally, synaptic vesicles alone do not demonstrate a synapse using electron microscopy. Rather, the presence of an active zone release site, seen as an electron dense area of the pre-synaptic plasma membrane in an electron microscopy micrograph, is what defines a synapse (Beaudoin et al., 2012;Burette et al., 2015). Bassoon is a well-characterized component localized to the active zone and found at synapses throughout the mammalian central nervous system (tom Dieck et al., 1998;Richter et al., 1999;Altrock et al., 2003).

Proteins localized post-synaptically come in one of two types. They can either be a neurotransmitter receptor or a post-synaptic scaffolding protein. Neurotransmitter receptors are localized at the postsynaptic membrane but may also be found on the plasma membrane away from the synapse and in intracellular vesicular stores (Bissen et al., 2019;Vieira et al., 2020). Postsynaptic scaffolding proteins are specifically localized to the postsynaptic membrane helping to anchor the neurotransmitter receptors along with the actin cytoskeleton and important signaling kinases and phosphatases necessary for inducing use-dependent long term changes in the synapses (Won et al., 2017;Groeneweg et al., 2018;Bissen et al., 2019;Pizzarelli et al., 2020).

The challenge is in identifying where the pre- and post-synaptic channels from the multi-labeled image overlap. The synapse is defined by the intersection of the two chosen markers, such as bassoon expression for the pre-synaptic compartment and a glutamate receptor for the post-synaptic compartment. Neither marker alone ideally identifies a synapse. Additionally, only including the overlap does not give a full depiction of the synapse.

A number of software tools for analyzing correlation, some of which are available free and open-source as add-ons to ImageJ, allow for identification of regions where two markers are overlapping (Zinchuk and Grossenbacher-Zinchuk, 2009;Singan et al., 2012;Lagache et al., 2015). Unfortunately, correlation analysis is not ideally suited for the imaging analysis required for identification of synapses. As the two markers are localized in different neurons, the two signals may be minimally overlapping if the synapse is being viewed perpendicularly. The two signals may by separated by as much as 200 nanometers depending on the marker pair (Dani et al., 2010). This distance is at the resolution of the fluorescent microscope, whether using widefield, confocal, or two-photon microscopy, due to the diffraction limit of light, meaning the two markers may be overlapping by only a few pixels. Super-resolution microscopy further increases the resolution, but this will further decrease the overlap in the signal between the two compartments (Dani et al., 2010;Beaudoin et al., 2012;Pennacchietti et al., 2017).

We have created a tool that automates the identification of synapses by identifying those puncta that overlap between the two markers. Our method is free and open-source, and combines the tools available as a part of ImageJ or FIJI (Schneider et al., 2012). Layered onto the identification of synapses through examining two images, the synapses can be further identified by overlap with cytoplasmic markers of either cell type. While the basic results from this analysis identifies the numbers of synapses to aid in identification of synaptic density, the results allow for detailed morphometric analysis of the total distribution of each marker individually and the distribution of synaptically localized markers. Finally, each individual puncta is matched to the overlapping puncta in the opposite channel. Thus, our method not only allows for automated detection of synapses, but further extends the analysis available to the user based upon the computed results. This analysis will allow for detection of changes in synaptic structure in addition to synaptic density.

## 2 Materials and Methods

### Animal care

Balb/c mice (Charles River Labs) were housed in microisolator cages with ad lib food and water. All procedures were approved by the Trinity University Animal Care and Use Committee.

### Stereotaxic surgery

Balb/c mice older than 3 months were used to inject AAV5 serotyped virus encoding CaMKII promoter driven expression of channelrhodopsin2 tagged with yellow fluorescent protein (ChR2-YFP) as previously described (Beaudoin et al., 2018). Briefly, mice were continuously anesthetized with 1-2% isoflurane delivered in 2 lpm oxygen. Lidocaine/Bupivacaine was used for nerve block at the incision site. Rimadyl was provided pre-, peri-, and post-operatively for pain management. Using a KOPF1900 stereotaxic frame, enabling precise correction of translation and rotation of the skull in all three axes, the pedunculopontine tegmental nucleus was bilaterally injected with 150 nL of AAV5-CaMKII-ChR2-YFP (~2×10^9^ vg/μL, UNC Vector Core) at the coordinates of −4.6 mm AP, −3.7 mm DV, and +/−1.2 mm ML. The virus injection rate was 50 nL/min, controlled by a syringe pump (WPI, UMP3) with a 5 minute wait period before withdrawing the injection needle (Hamilton 7000.5 syringe). The scalp was closed with VetBond. Animals were monitored for signs of distress, with all animals showing excellent recovery within a few days.

### Perfusion of mice

Mice were anesthetized with isoflurane and then intraperitoneally injected with 15 μL/g of 2.5% tribromoethanol in tert-amyl alcohol diluted in sterile saline. Mice were perfused through their aorta with 10 mL of ice-cold PBS, pH=7.4, followed by 25 mL of ice-cold 4% paraformaldehyde in PBS, pH=7.4. The dissected brain was post-fixed for 2 hours in 4% paraformaldehyde, and then cryoprotected with an ascending series of sucrose solutions in PBS, 10%, 20%, and 30%, incubating the brain overnight in each solution. The brain was then blocked and mixed in freezing medium (OCT) before freezing on dry ice.

### Sectioning and Immunostaining of Tissue

Coronal sections were prepared on a cryostat at 20 μm and immediately mounted on slides. Slides were maintained at −80 °C until ready for staining. Every 10^th^ section was Nissl stained with Cresyl Violet to identify sections corresponding to areas of interest (Franklin and Paxinos, 2013).

Immunostaining was done as described previously (Beaudoin et al., 2012). The following primary antibodies were used mouse anti-bassoon (1:400, ABCAM ab82958), rabbit anti-gephyrin (1:1000, ABCAM ab32206), rabbit anti-NMDA receptor subunit 1 (1:1000, ThermoFisher PA3-102), and chicken anti-tyrosine hydroxylase (1:500, ABCAM ab76442). Secondary antibodies used were goat anti-mouse (1:1000 for all; Alexa568, ABCAM ab175473; Alexa488, Fisher A11029; Alexa647, Fisher A21241), goat anti-rabbit (1:1000 for all; Alexa488 ABCAM ab150077; Alexa568, Fisher A11036), and/or goat anti-chicken (1:1000, Alexa405, ABCAM ab175675). Coverslips were mounted with ProLong Gold antifade (Invitrogen).

### Imaging and Analysis

All images were acquired on a Nikon A1R+ confocal equipped with laser lines at 405, 488, 563, and 647nm. Images were acquired at Nyquist resolution using either a 100x oil objective or a 60x oil objective. Laser power and PMT gain and offset were set to maximize use of the full range of intensity values with pixel intensities below the maximal intensity of 4095 for the synaptic channels. Multichannel images were acquired sequentially to minimize collection of closely overlapping emission ranges.

All analyses were done in batch including a no primary control slide to ensure the fidelity of the analysis software. Analysis settings and output are detailed below.

## 3 Results

### Overview of macro function

Putative synapses are identified via determining overlapping pre- and post-synaptic puncta. The macro first identifies the pre-synaptic puncta from one color channel and the post-synaptic puncta from a second color channel from the same image (Fig. 1A). This initial identification is key, and we have identified two methods of identification based upon the density of staining. For sparsely labeled images, a simple threshold of intensity values, above which is all considered signal and below which is all considered background will work (Fig. 1A, B). This requires that puncta are largely discrete and non-adjacent, surrounded by low intensity borders. In many ways, this type of puncta identification is both routine and well appreciated as a cornerstone of ImageJ analysis of features (Beaudoin et al., 2012;Dzyubenko et al., 2016;Timothy and Forlano, 2019;Taylor et al., 2020;Gautier and Ginsberg, 2021).

**Figure 1.**
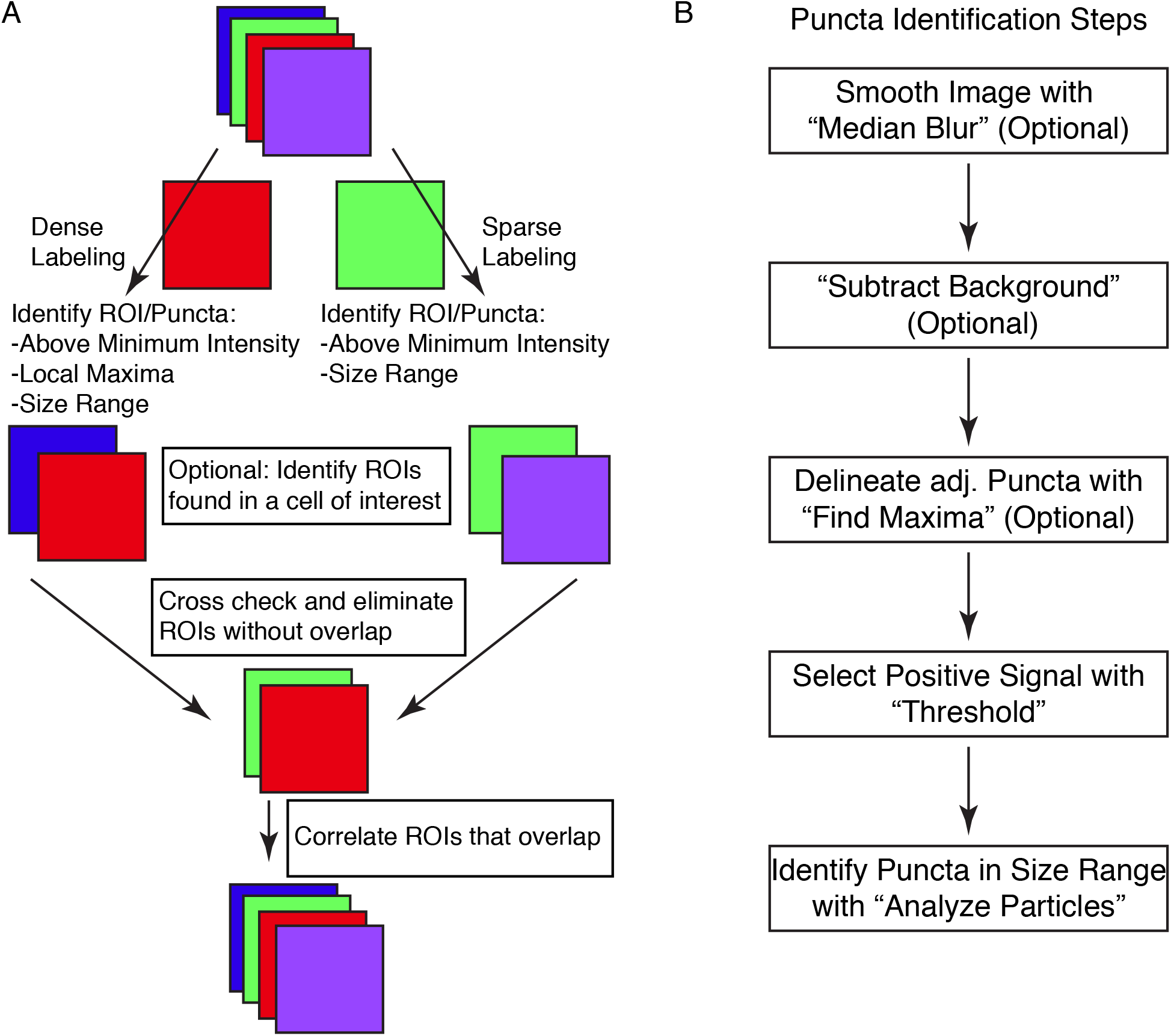
SynapseJ identifies putative synaptic contacts through identification of puncta in single channels and then examines overlap between channels. (**A)** Initial analysis of pre- or post-synaptic puncta is dictated by the density of puncta in the image. Before identifying correlation between pre- and post-synaptic puncta correlated, one or both channels may be selected for overlap with a cell-specific marker. The final multi-channel image recombines data from each channel, using only the processed image. **(B)** The identification of puncta proceeds through only a few steps. Smoothing the image with “Median Blur” removes shot noise in the image. A non-uniform hazy background is removed with “Subtract Background”. For densely labeled images, adjacent puncta are delineated with “Find Maxima”. Ultimately, densely and sparsely labeled images are analyzed for puncta by selecting a “Threshold” intensity for positive signal, followed by “Analyze Particles” to remove puncta not within the appropriate size range. Once completed, a list of puncta are identified for that single channel.

For densely labeled images, in which the puncta are not discrete and are adjacent, we have devised a second method of puncta identification. The second method of puncta identification is by combining the functionalities of thresholding with a local maxima identification function of ImageJ titled “Find Maxima” (Fig. 1A, B). This built-in function of ImageJ can identify local maxima ignoring those that appear below the threshold. While many users have found this works well for identifying points of interest, another utility of this function is to identify the borders between adjacent puncta, that are not as clearly delineated. This function is enabled by selecting the “Segmented particles” option. As this functionality can not be applied to Z-stack images, we have created a stand alone function that applies the segmented particles option of the “Find Maxima” function to each individual slice of the Z-stack. Using one of these two puncta identification methods, the macro stores each puncta as part of the ROI manager. This list is saved so the user can refer back to the originally identified puncta. Examination of the list is key in being sure that the puncta are not missed or merged.

The two lists of puncta are then compared for overlap using a graphical approach (Fig. 1A). Each list of puncta is checked for overlap with puncta in the opposite channel. This is done as a brute force computation, with each puncta checked and either saved or deleted. Once both lists of puncta have been culled for non-overlapping puncta, the macro then sets to identify which punctum overlaps the punctum(a) of the other channel. In some cases, a single punctum from one channel may overlap with more than one punctum of the other channel. This data is used to create the correlation analysis tables which are listed both as pre-synaptic puncta vs. each overlapping post-synaptic puncta and as post-synaptic puncta vs. each overlapping pre-synaptic puncta.

All of the analysis is performed both quantitatively and graphically. The morphometry of the puncta before and after culling for overlap with the other channel are output. This information is collated together as a single tab-delimited file as well as individual files for each channel for each image. The graphical output means that the images are updated throughout the process. This includes identification of the original list of puncta and then after identification of overlapping puncta. The macro merges these processed images back with any other remaining channels and saves the final image as a TIFF file.

To run the macro, the user is prompted for information three times. First, the user is asked for the directory containing the images to be analyzed. Currently, the macro can use multichannel TIFF images (ending in .tif) or multichannel Nikon confocal images (ending in .ND2). These endings can easily be swapped out in the code, or files can be converted into a TIFF file. Second, the user is prompted for a directory to store the output from the macro. This should be a new folder, given the large amount of information to be generated. The final prompt is for the settings to be applied to analyze the images.

The macro is run in batch mode in ImageJ, which means that the only output to the screen will be to text windows. Initially, a log window is created to serve as a record of the settings being used for analysis and for monitoring macro progress, image by image. The results at intermediate stages are all saved. Then, after the macro is finished the analysis for each image is saved. Finally, once all the images are analyzed, a summary table listing the numbers of identified puncta from each channel from each image is output to a table and saved.

To demonstrate the effectiveness of this approach, we have applied the macro to analysis of immunostaining for the presynaptic scaffolding protein bassoon and the postsynaptic scaffolding protein gephyrin. Bassoon indiscriminately labels most synapses, being found at synapses for excitatory, inhibitory and modulatory neurotransmitters (tom Dieck et al., 1998;Richter et al., 1999;Cui et al., 2021). In contrast, gephyrin is found at inhibitory synapses comprised of either GABA or glycine receptors (Craig et al., 1996;Pennacchietti et al., 2017;Groeneweg et al., 2018;Pizzarelli et al., 2020). Gephyrin is thought to be found at a subset of inhibitory synapses (Groeneweg et al., 2018;Pizzarelli et al., 2020). Thus, immunostaining for bassoon will be dense and adjacent without clear borders, while gephyrin is comparatively sparse and only minimally adjacent and has clear borders. Systematic variation of the analysis parameters identifies the limits of this analysis. We examined synaptic labeling in ventromedial hypothalamus (VMH), central amygdala (CeA), basolateral amygdala (BLA), and the arcuate (Arc). We also examined the effects of adjusting the parameters for analysis on the number of bassoon and gephyrin puncta in each brain region, using staining in BLA for illustrative purposes.

### Identification of Densely-labeled Puncta

For densely-labeled puncta identification, there are three variables that affect their identification, which are the noise tolerance or prominence value for the “Find Maxima” function, the threshold intensity, and the size of individual puncta (Fig. 1B). The images are smoothed before attempting to identify puncta to remove noise introduced by using a point scanning confocal microscope with a photomultiplier tube for assigning fluorescent intensity values (Fig. 1B). To smooth the image, the macro can optionally apply a median blur (Fig. 2A, D). A sub-pixel radius does not affect the puncta analysis, but does lead to visual improvement of the image (Fig. 2A, D).

**Figure 2.**
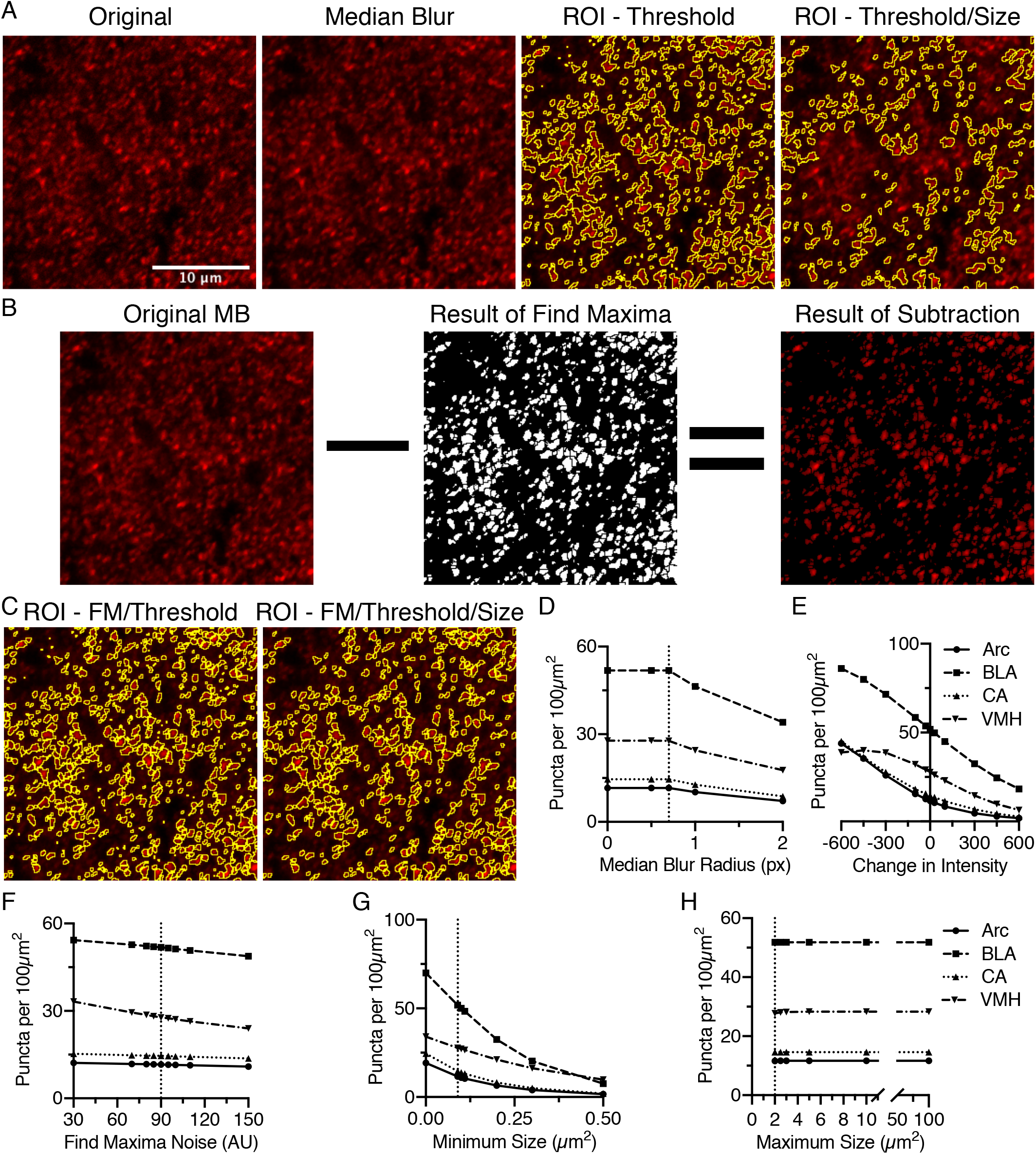
Identification of puncta in densely labeled images requires a number of settings, but is most affected by the threshold intensity and the minimum size. **(A)** Anti-bassoon staining in BLA is densely labeled but clearly denotes puncta. Median blur with a sub-pixel radius removes noise without noticeably affecting the puncta. Identifying puncta using analyze particles above a minimum threshold, removing small and large puncta, leads to significant loss of most of the stained puncta. **(B)** Alternatively, the “Find Maxima” function (FM) identifies borders between puncta above the threshold using the segmented particles setting. This creates a black and white image, which with “Image Math”, can be used to remove pixels below threshold and create borders around closely spaced puncta. **(C)** Anti-bassoon puncta identified using thresholding and FM is further refined by using the analyze particles function to remove puncta outside a minimum and maximum size range (Size). Number of puncta identified in anti-bassoon staining in four brain regions is maximally, and universally negatively correlated with changes in intensity threshold **(E)** and a minimum size **(G)**. Sub-pixel median blur **(D)**, changes in the noise setting for FM **(F)**, and maximum size **(H)** have comparatively little effect on the number of identified puncta. Vertical dotted lines denote the original starting setting used for analysis. Scale bar denoted in A is 10μm for all images.

The initial key parameter is the threshold intensity. The threshold intensity should be set to identify the puncta at their brightest, not merely that they are above background. Even with very narrow optical section thickness, puncta can be seen to increase in intensity as they are imaged in the correct focal depth (Fig. 2A). Even with a high threshold intensity, in densely labeled areas, many puncta will be adjacent to each other without clear borders (Fig. 2A). This identifies the key issue of solely using threshold to identify puncta, as the puncta will be merged into large regions of interest (ROIs), and then eliminated as being too large (Fig. 2A). This parameter is one of the most critical in identifying puncta. We found a negative correlation between threshold and puncta density, with far more puncta identified with lower thresholds and fewer puncta identified at higher thresholds (Fig. 2E). In most instances, this trend was super linear, although in one instance at lower thresholds the trend became sub-linear (Fig. 2E).

Adjacent puncta are distinguished by the “Find Maxima” function which examines the image for local maxima and based upon the decrease in intensity between two maxima in the two-dimensional image distinguishes the border between the two images (Fig. 2B). This function requires limiting the analysis to above the intensity threshold and the segmented particles options. The tuning of this function is through modulating the noise tolerance, in ImageJ, or prominence, in FIJI. This parameter indicates the difference in intensity required to distinguish small fluctuations in intensity from the start of the next local maxima. The preview function is instrumental in real time feedback required to distinguish the value that ideally identifies individual puncta versus noise. The output is a black and white image distinguishing the extent of each puncta, creating a one-pixel wide border between adjacent puncta. As this image discards the intensity values from the original image, image math is used to remove pixels below threshold and introduce a border between puncta (Fig. 2B, C). Changes in the noise tolerance for the “Find Maxima” function were inversely correlated with the number of identified puncta (Fig. 2F). This relationship was relatively linear (Fig. 2F). Surprisingly, the effects on puncta density are relatively minimal (Fig. 2F).

The final consideration for identification of synaptic puncta is size, both the minimum and maximum allowable sizes. The range of acceptable sizes should be set empirically based upon your particular region of analysis. For the ideal minimal size, one must consider the constraints of the imaging method. Nyquist sampling on a confocal with a high NA, high magnification objective, the square pixels are between 0.08-0.1 μm on an edge. However, given the diffraction limit of light and a pixel size of 0.1 μm^2^, the minimum size would be 3-4 pixels by 3-4 pixels corresponding to an area of 0.09-0.1 μm^2^. With a minimum size of 0.09 μm^2^, the puncta identified by threshold and the “Find Maxima” function are further refined (Fig. 2C, G). The minimum size can lead to dramatic shifts in measured puncta density (Fig. 2G).

The maximum size should be set to exclude cell body and dendritic staining. Generally, the maximal size would be less than 5 μm^2^, but may be no larger than 2 μm^2^. The maximal size depends on the known types of synapses in the specific brain region being. In all four areas tested, increasing the maximal size from 2 μm^2^ had minimal effects on puncta density (Fig. 2H).

### Identification of Sparsely labeled Puncta

For sparsely labeled puncta, the macro uses two parameters to identify the relevant puncta, including the intensity threshold and size of puncta (Fig. 1B). Optionally, the channel may be smoothed using a median blur, and based upon the analysis of densely-labeled puncta, a one pixel radius or smaller would seem appropriate (Fig. 1B and 3A). Sparsely labeled puncta are easily distinguished above background and are delineated graphically by thresholding the image (Fig. 3B). While this delineates individual puncta, some of these may still be too small or large to correspond to relevant staining. Thus, the puncta are qualified by rejecting puncta that are too small or too large. The range of acceptable sizes should be set empirically based upon your particular region of analysis, but follow the same size constraints as detailed above (Fig. 3B).

**Figure 3.**
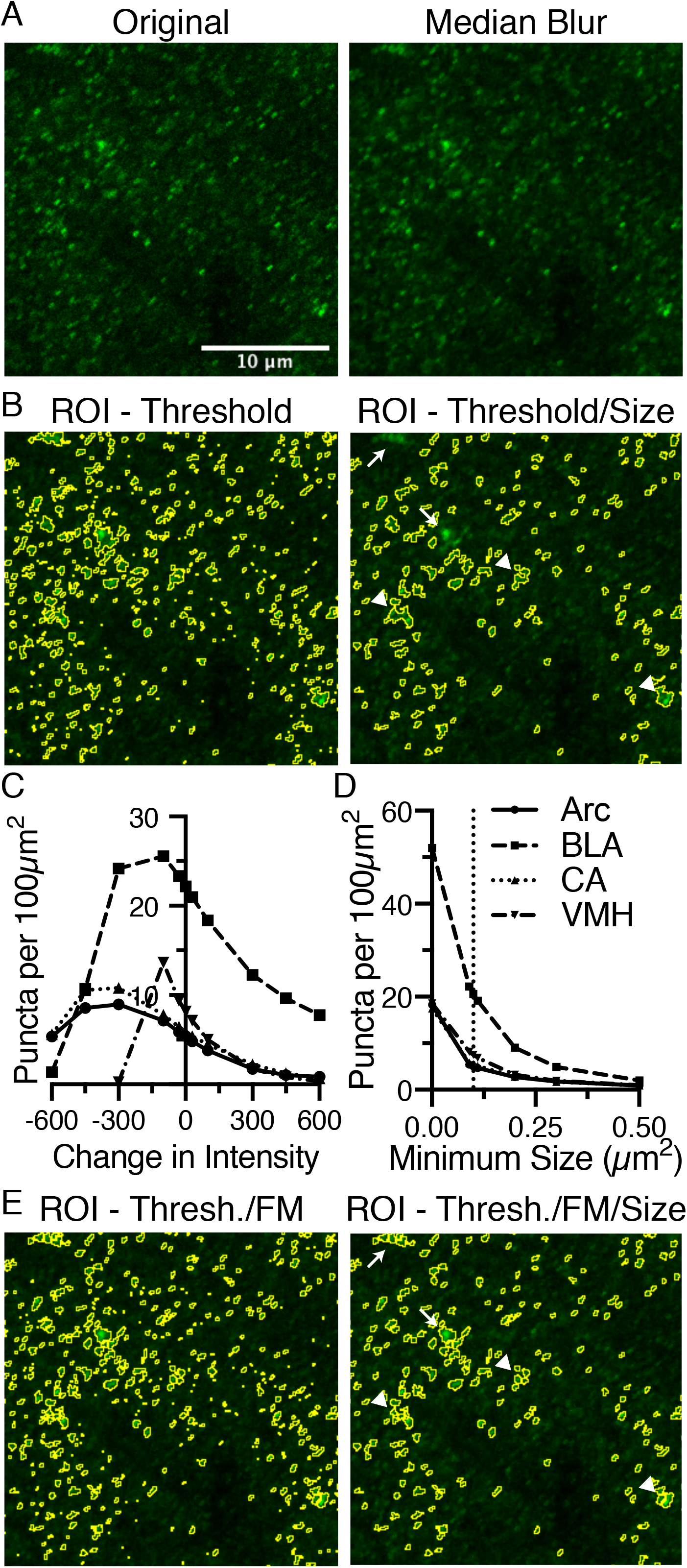
Identification of puncta in sparsely labeled images are dictated by an intensity threshold and minimum size and may still be enhanced by using the same process as densely labeled images. **(A)** Anti-gephyrin staining in BLA is one such marker that is sparsely labeled. The original image is median blurred to smooth noise. **(B)** The ROIs, circled in yellow, selected based upon a threshold intensity are further refined by removing puncta that are too small or big. **(C)** Gephyrin puncta was examined in four brain regions (Arc, dot continuous line; BLA, squares and dashed line; CeA, triangles and dotted line; VMH, inverted triangle and dash-dot line) Puncta number are negatively correlated with changes in threshold intensity, until the threshold has been reduced too far. **(D)** Puncta number is also negatively correlated with changes in minimum size. **(E)** Reanalysis of anti-gephyrin staining in BLA using the dense analysis method identified different puncta. Puncta denoted by arrows are absent from the final image in B while present in the final image in E. Puncta denoted by arrows are present in image B, but have been sub-divided in E. Scale bar denoted in A is 10μm for all images.

In varying these two parameters, the threshold and the range of acceptable sizes, we found two very different trends in puncta density (Fig. 3C, D). Varying the threshold was negatively correlated with puncta density in all four brain regions (Fig. 3C). However, as the intensity threshold was lowered further, the puncta density then decreased. This decrease in puncta density at the lowest intensity thresholds will be addressed later in the results section.

In the specific example of gephyrin puncta analysis in BLA shown in figure 3, some puncta clusters are merged into a single larger punctum, and in some cases these larger puncta are removed due to the maximum size (Fig. 3B). Thus, despite the sparseness of gephyrin puncta, perhaps this channel would also benefit from the “Find Maxima” approach to identifying puncta. Using this alternate approach, these local clusters of gephyrin are successfully subdivided into smaller puncta (Fig. 3E). These subdivided clusters lead to inclusion of formally excluded puncta, as they are now within the size threshold (Fig. 3E). Thus, the densely-labeled puncta identification method is more generally applicable to all densities of immunostaining.

### Identification of overlapping puncta

Once puncta have been identified in either channel, the macro eliminates the puncta that do not overlap with each other (Fig. 4A). The remaining, overlapping puncta are the putative synapses, which are inhibitory in the case of gephyrin staining. In some cases, this will lead to a dramatic reduction in puncta density, as is the case for bassoon puncta in VMH, starting with a puncta density of ~30 puncta per 100 μm^2^ and ending with ~3 puncta per μm^2^ (Fig. 2 and 4). We have analyzed the effects of changes in initial identification of puncta affect the identification of these putative synaptic puncta, in both channels. This analysis includes how modifying identification of bassoon puncta affects the resulting gephyrin-positive bassoon puncta, and the bassoon-positive gephyrin puncta (Fig. 4B-C). In this case, the identification of gephyrin puncta was the same so the starting number of gephyrin puncta was not altered and thus not plotted.

**Figure 4.**
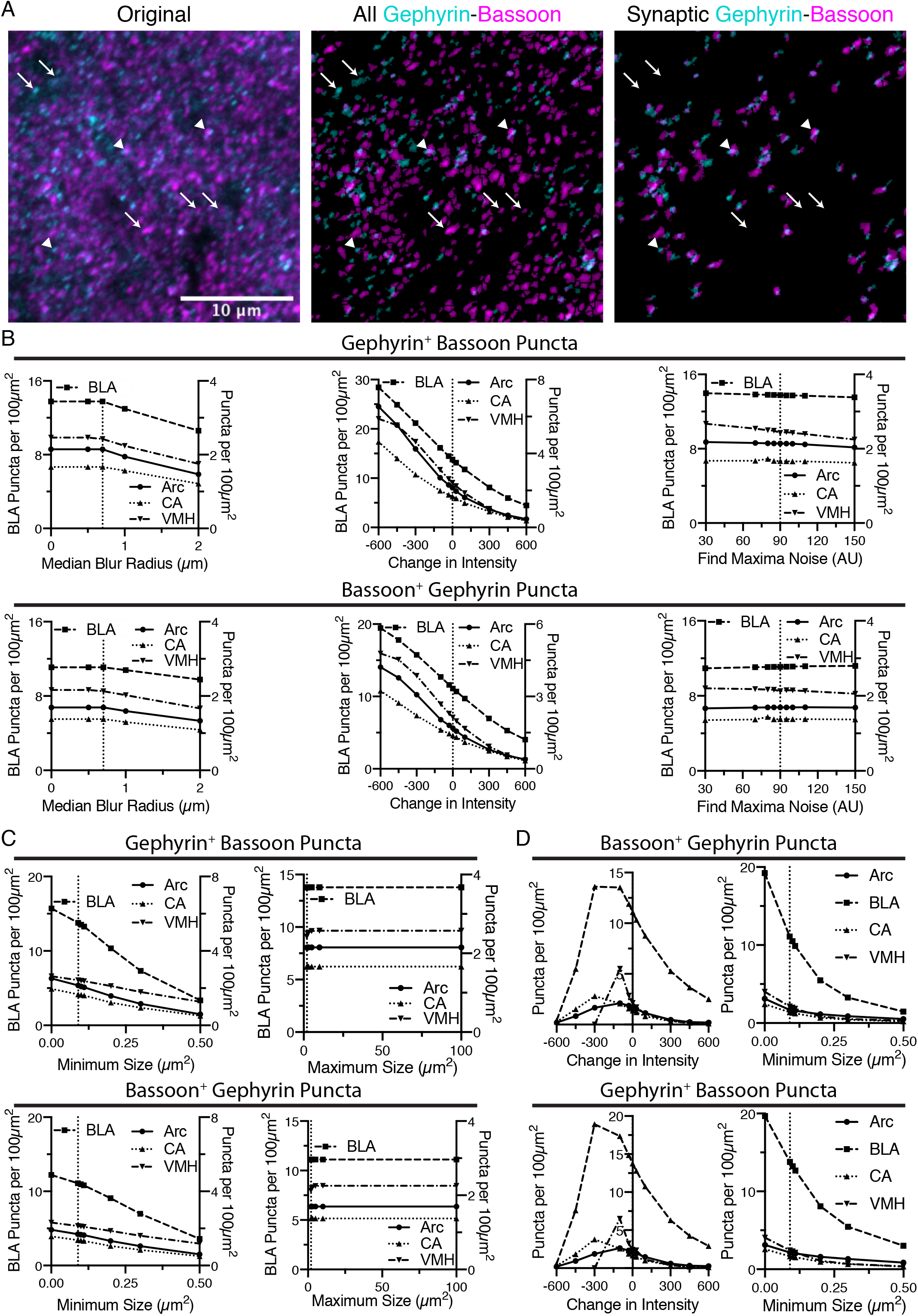
Identification of putative synapses is done by finding overlap between puncta identified in the two images. **(A)** In the following sequence of images, gephyrin is colored in cyan and bassoon colored in magenta. Shown is the original two channel image, all of the identified puncta in both channels, and then the final identification of overlapping puncta. Some puncta are eliminated (identified with white arrows), while remaining, overlapping puncta are putative synaptic contacts (identified with white arrowheads). Scale bar,10 μm, applies to all images. **(B)** Increasing the threshold intensity for bassoon puncta reduces synaptic puncta density, for either gephyrin-positive bassoon puncta or bassoon-positive gephyrin puncta. Increasing radius for median blur or the noise value for “Find Maxima” cause relatively little changes in synaptic puncta density. Trends in synaptic puncta density are largely similar between brain regions. **(C)** Increasing minimum size of synaptic puncta decreases puncta density across all brain regions, while increasing maximum size leads to little change in synaptic puncta density. **(D)** Similar to the dense puncta identification, using the sparse identification method leads to synaptic puncta being negatively correlated with changes in minimum size of puncta. However, decreasing the intensity threshold too much leads to a decrease in synaptic puncta density.

In general, the parameters that led to changes in identification of bassoon puncta, similarly altered the putative synaptic puncta. The median blur radii of 1 pixel or less leads to little change in the number of gephyrin-positive bassoon puncta or bassoon-positive gephyrin puncta (Fig. 4B). Similarly, modulating the noise tolerance/prominence value for the “Find Maxima” function (Fig. 4B) or the maximum size of puncta (Fig. 4C) had little effect on the puncta density of gephyrin-positive bassoon puncta or bassoon-positive gephyrin puncta. However, both changes in threshold intensity (Fig. 4B) and minimum size (Fig. 4C) leads to dramatic changes in identification of gephyrin-positive bassoon puncta or bassoon-positive gephyrin puncta. It is notable however, the dramatic difference in slope for the minimum size between initial identification of bassoon puncta (Fig. 2G) and the subsequent identification of gephyrin-positive bassoon puncta or bassoon-positive gephyrin puncta (Fig. 4C), in particular for removing the minimum size. This suggests that many of the smallest bassoon puncta are eventually eliminated as they do not overlap with gephyrin.

We also examined the effects of changing intensity threshold and minimum size for gephyrin puncta on identification of bassoon-positive gephyrin puncta or gephyrin-positive bassoon puncta (Fig. 4D). In general, the trends seen for identification of gephyrin puncta (Fig. 3 C, D) resemble the trends for bassoon-positive gephyrin puncta or gephyrin-positive bassoon puncta (Fig. 4D). Similar to bassoon staining, removal of a gephyrin minimum size caused a greater than two-fold increase in gephyrin puncta (Fig. 3D), but are less than two-fold increase in bassoon-positive gephyrin puncta or gephyrin-positive bassoon puncta (Fig. 4D). Thus, the macro appears to identify overlapping puncta, with some of the same constraints as the initial identification of puncta.

### Synaptic Analysis from the data

Thus far, the data analyzed has been for puncta density as a readout. A key feature of this analysis is the detailed morphometry data provided as output. One can examine the mean size, intensity, and integrated density of both Bassoon and Gephyrin puncta, before and after qualifying the puncta for overlap. Given the large changes in puncta density induced by changing the threshold intensity or minimum puncta size of immunostaining for bassoon or gephyrin, we focused this more detailed analysis on these parameters (Figs. 5, 6). Increasing the threshold intensity for bassoon or gephyrin increased the mean intensity of bassoon or gephyrin puncta (Fig. 5A, D). Increasing the threshold intensity similarly increased gephyrin-positive bassoon and bassoon-positive gephyrin (Fig. 5B, E). However, increasing the threshold intensity for bassoon or gephyrin had no effect on bassoon-positive gephyrin or gephyrin-positive bassoon, respectively (Fig. 5C, F). In general, increasing the threshold intensity caused a decrease in area of bassoon puncta, independent of gephyrin, and gephyrin, independent of bassoon (Fig. 5A, B, D, E). The effects of these two changes in integrated density were unpredictable, with generally flat or decreasing integrated density of any type of bassoon puncta when identifying bassoon puncta (Fig. 5A, B), but increasing or flat integrated density for all types of gephyrin puncta when identifying gephyrin puncta (Fig. 5D, E). Interestingly, increasing the threshold intensity for bassoon had negligible effects on area or integrated density of bassoon-positive gephyrin puncta from BLA, CeA, and VMH, but increased the size and integrated density of bassoon-positive gephyrin puncta in Arc (Fig. 5C). However, increasing the threshold intensity for gephyrin increased the size and integrated density of gephyrin-positive bassoon puncta from all four brain regions (Fig. 5F). Two interesting points should be noted. First, changing the threshold intensity reduces the size and increases the mean intensity, with minimal changes on integrated density of identified puncta. Second, while the changes in raw identified puncta are correlated with overlapping puncta from the same imaging channel, the overlapping puncta from the other channel have limited changes to area of puncta, with no effect on mean intensity.

**Figure 5.**
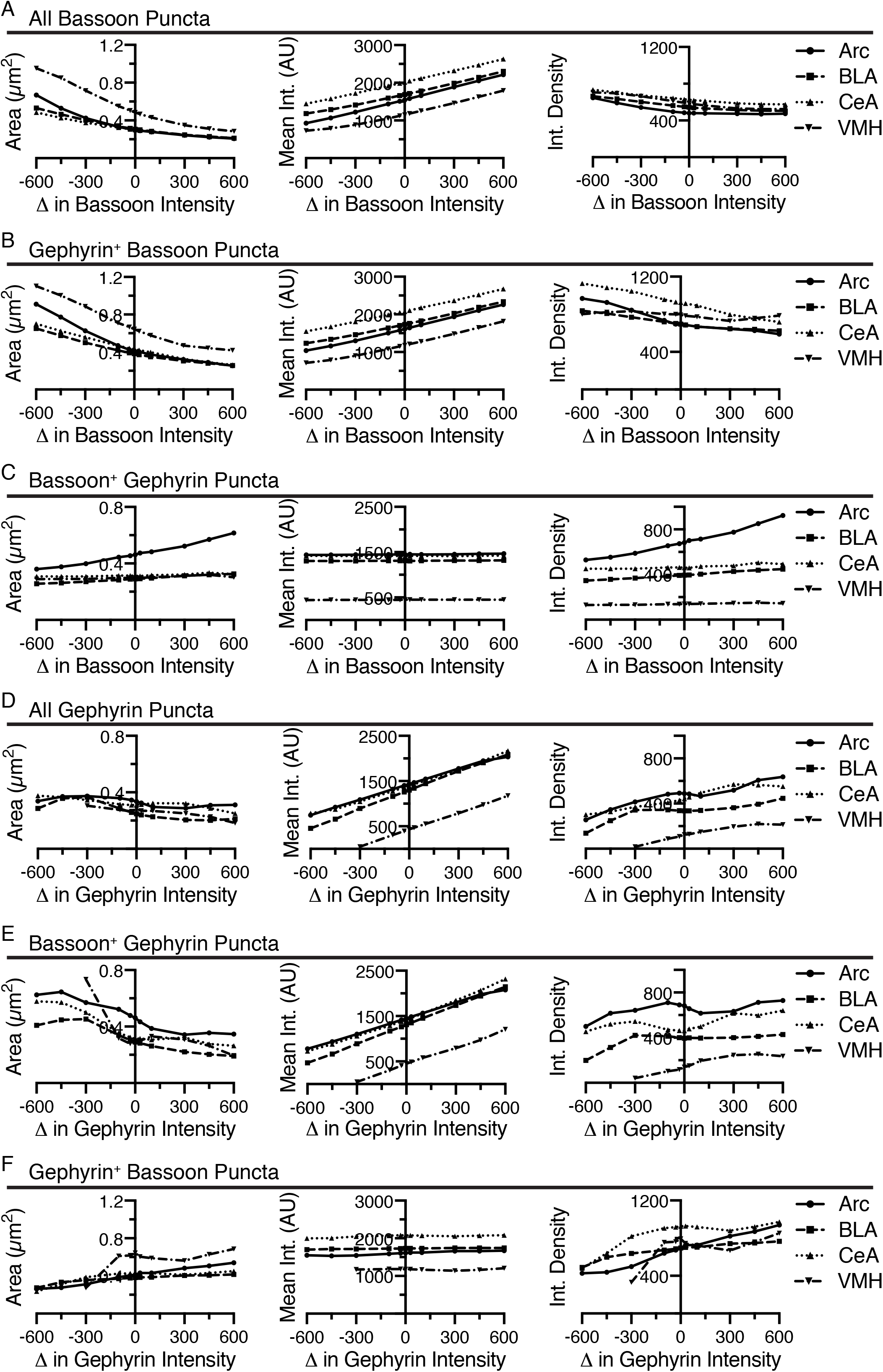
Increasing the threshold intensity for identification of bassoon or gephyrin, generally decreased area, increased mean intensity, and decreased integrated density for bassoon and gephyrin puncta, respectively, with inconsistent effects on associated gephyrin or bassoon puncta. Changes in size (area, μm^2^), average intensity, and integrated density of all bassoon puncta **(A)**, gephyrin-positive bassoon puncta **(B)**, and bassoon-positive gephyrin puncta **(C)** are plotted versus changes in threshold intensity for identification of bassoon puncta. Similarly, changes in size (area, μm^2^), average intensity, and integrated density of all gephyrin puncta **(D)**, bassoon-positive gephyrin puncta **(E)**, and gephyrin-positive bassoon puncta **(F)** are plotted versus changes in threshold intensity for identification of gephyrin puncta.

**Figure 6.**
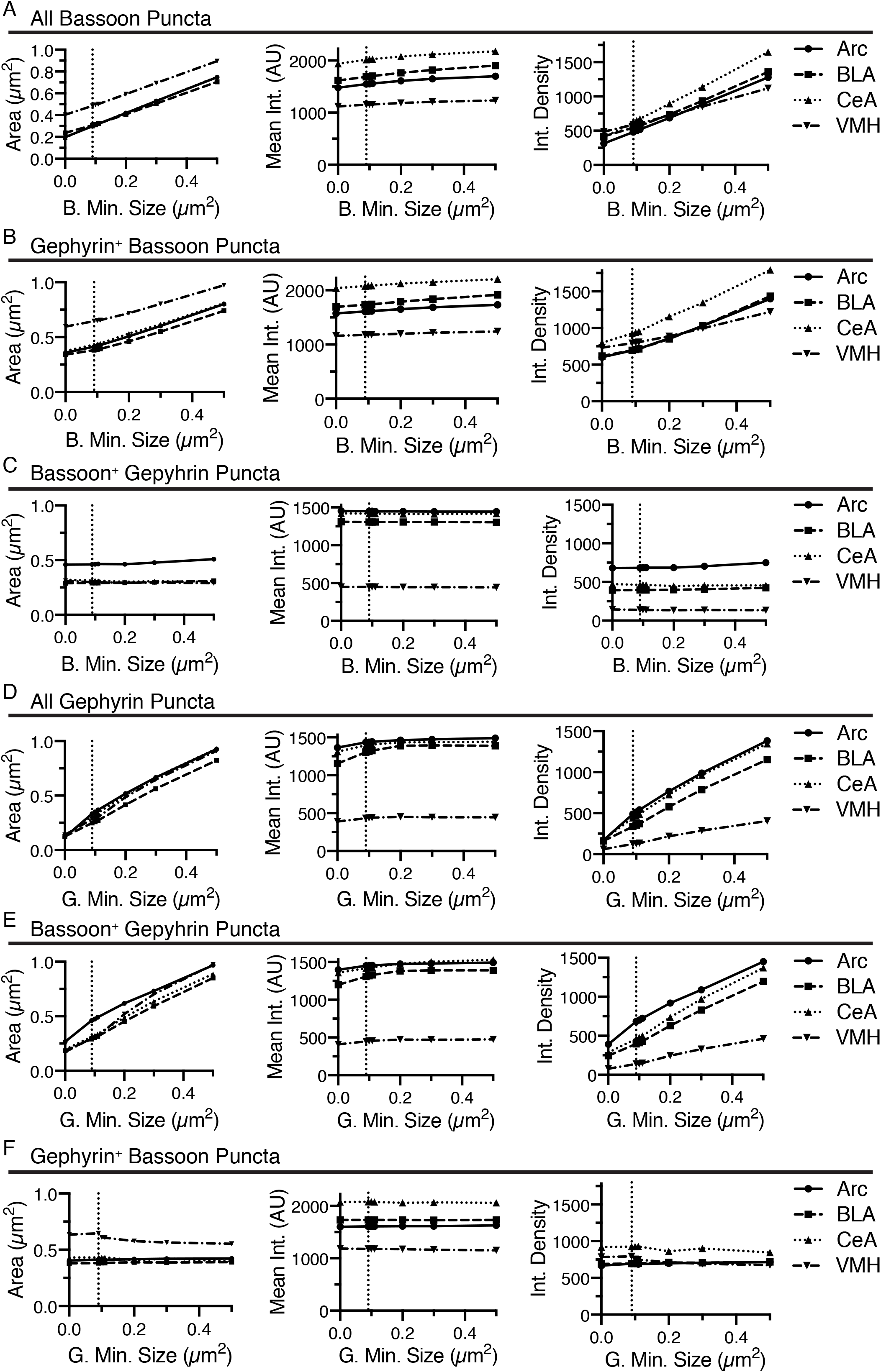
Increasing the minimum size for identification of bassoon or gephyrin, generally increased area, slightly increased mean intensity, and increased integrated density for bassoon and gephyrin puncta, respectively, with inconsistent effects on associated gephyrin or bassoon puncta. Changes in size (area, μm^2^), average intensity, and integrated density of all bassoon puncta **(A)**, gephyrin-positive bassoon puncta **(B)**, and bassoon-positive gephyrin puncta **(C)** are plotted versus changes in minimum size for identification of bassoon puncta. Similarly, changes in size (area, μm^2^), average intensity, and integrated density of all gephyrin puncta **(D)**, bassoon-positive gephyrin puncta **(E)**, and gephyrin-positive bassoon puncta **(F)** are plotted versus changes in minimum size for identification of gephyrin puncta.

Increasing the minimum puncta size for analysis of bassoon or gephyrin immunostaining induced increases in area of bassoon and gephyrin puncta, independent of overlap with the other channel (Fig. 6A, B, D, E). While increasing the minimum size of bassoon and gephyrin puncta had little effect on the mean intensity of bassoon or gephyrin puncta, regardless of overlap with the alternate channel, respectively, the integrated density of these same puncta increased, due to the changes in size (Fig. 6A, B, D, E). Modulating the minimum puncta size of bassoon or gephyrin had little effect on size, mean intensity, or integrated density for gephyrin-positive bassoon or gephyrin positive-bassoon, respectively (Fig. 6C, F).

One question easily answered by this analysis is how the puncta classified by their overlap with the other channel compares to the original list of puncta. Comparison of the puncta density has already highlighted the significant reduction in puncta density when selecting for the overlapping puncta. There are also upward shifts in average size, mean intensity and integrated density that can be noted when comparing all bassoon (Figs. 5A and 6A) to gephyrin-positive bassoon (Figs. 5B and 6B). For gephyrin puncta, these effects appear to be most dramatic for Arc and much more subtle for BLA, CeA, and VMH (Figs. 5C, D and 6C, D). Using BLA as an example, we created histograms of the fraction of puncta at a range of areas, mean intensities, and integrated densities to directly compare the collection of all bassoon to gephyrin-positive bassoon (Fig. 7A) and gephyrin to bassoon-positive gephyrin puncta (Fig. 7B). For bassoon puncta in BLA, the prevalence of smaller puncta is reduced while larger puncta are more prevalent in the synaptic fraction. Additionally, the mean intensity of gephyrin-positive puncta is shifted to higher values, which in combination leads to a noticeable increase in integrated density (Fig. 7A). Bassoon-positive gephyrin shows no shift in size or integrated density, but has a noticeable change in mean intensity of puncta (Fig. 7B). The original distribution of gephyrin intensities appears bimodal, while the synaptic gephyrin puncta appear unimodal (Fig. 7B).

**Figure 7.**
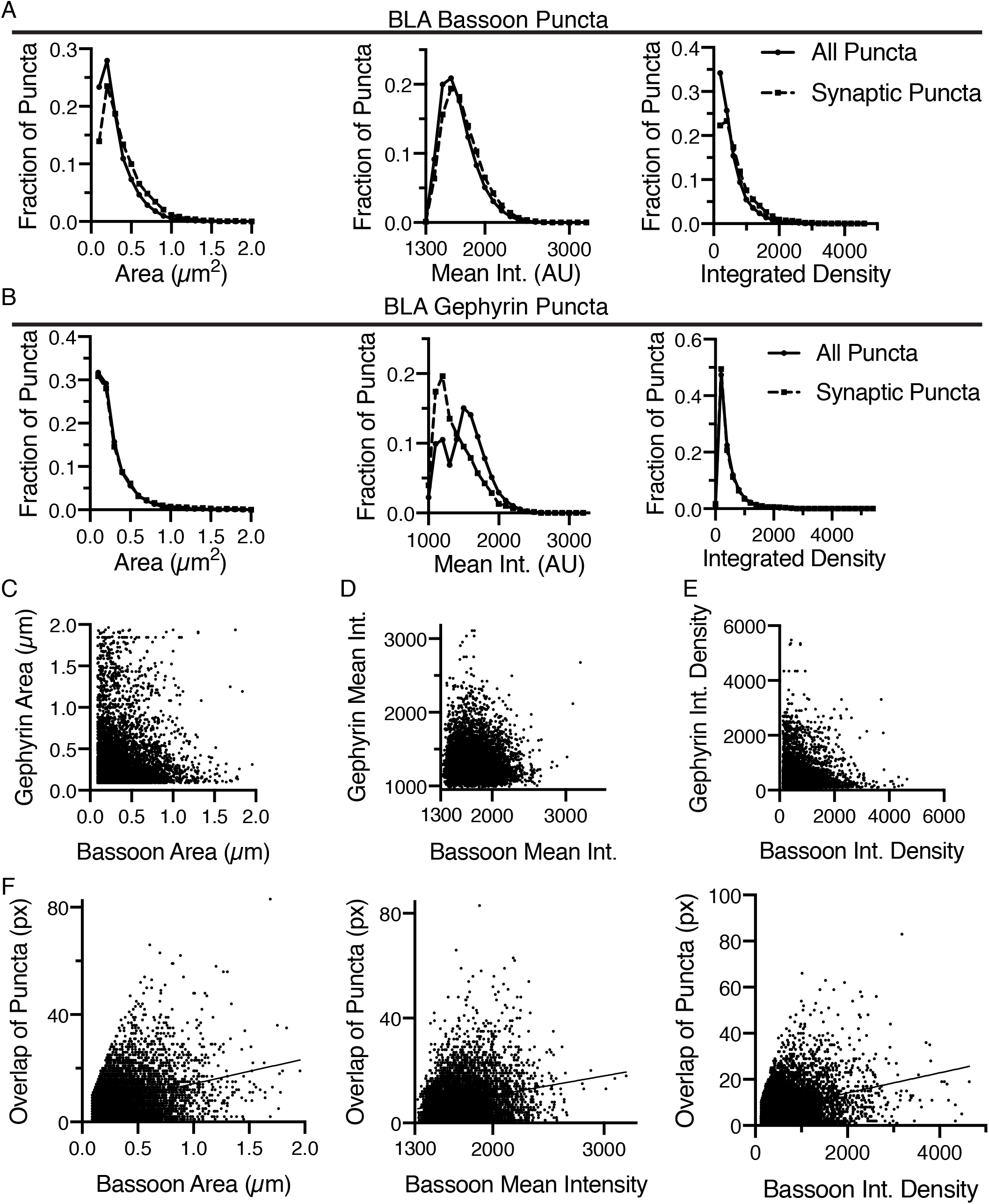
Identification of overlapping bassoon and gephyrin puncta in the BLA enables detailed morphometric analysis. **(A)** In the BLA, the distribution of area, mean intensity, and integrated density of all bassoon puncta and synaptic bassoon puncta reveals changes in distribution. **(B)** Similarly, the distribution of area, mean intensity, and integrated density of all gephyrin puncta and synaptic gephyrin puncta in the BLA reveals changes in distribution. Identification of overlapping bassoon and gephyrin puncta enable correlation of area **(C)**, mean intensity **(D)**, and integrated density **(E)**. **(F)** Overlap between gephyrin and bassoon puncta is positively correlated with area, mean intensity and integrated density of bassoon puncta in BLA.

Another key feature of the analysis is the matching of Bassoon puncta to individual Gephyrin puncta and vice versa. Many regions of the nervous system have different inhibitory inputs, which may create inhibitory synapses of different sizes or types. This is information that may be discerned through this analysis. For instance, the analysis allows for examining correlation of pre- and post-synaptic puncta size, intensity, and integrated density (Fig. 7C-E). The software also computes the distance between the geometric centers of each pre-and post-synaptic puncta, as well as between the intensity center of each puncta. Finally, the software counts the number of pixels corresponding to the overlap between the puncta. Thus, we can correlate the area, intensity, and integrated density of individual puncta with the amount of overlap (Fig. 7F). This showed a positive correlation between the amount of overlap and the area, mean intensity and integrated density of gephyrin-positive bassoon puncta (Fig. 7F). This detailed analysis may allow for identification of characteristics of different classifications of synapses.

### Additional image processing parameters

To address the concern that identified puncta need only overlap by a single pixel might lead to a number of false positive overlapping puncta, one option would be to erode the perimeter of the puncta by a single pixel. By reducing the puncta size, and then examining the overlap, only those puncta that are most apposed to each other would remain. However, this would lead to a significant decrease in the number of puncta, and may have non-linear effects on puncta based on original size. Instead, to reduce the number of potential false positive synapses, the number of pixel overlap can be specified (Fig. 8A). Shown is the overlapping bassoon-gephyrin puncta in BLA as identified in figure 4A, now limiting to only those puncta that overlap by 6 or more pixels (Fig. 8A). The size of the puncta may alternatively be decreased or eroded and increased or dilated to attempt to address false positive or negative overlapping puncta.

**Figure 8.**
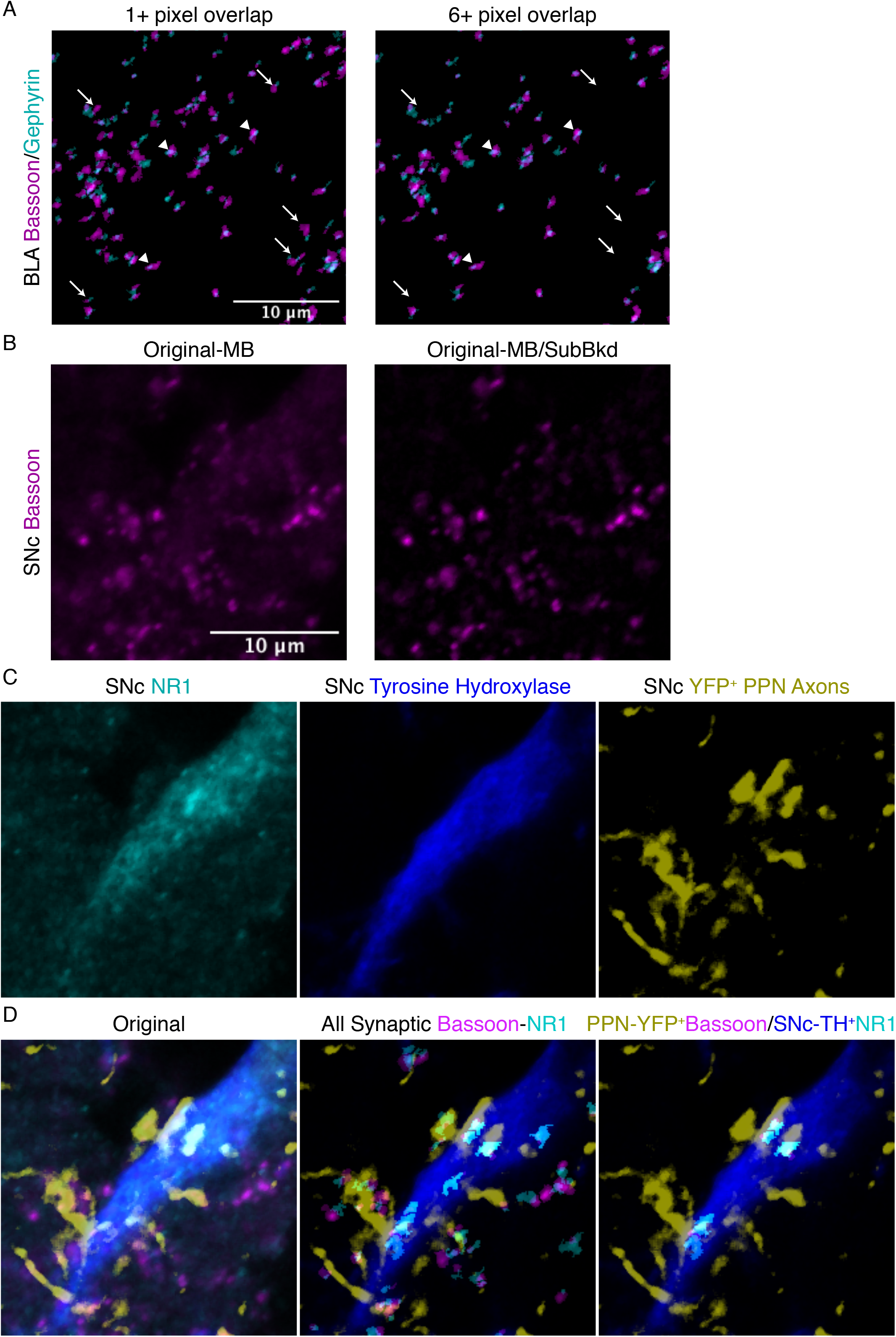
Additional features of SynapseJ allow for identification of synaptic puncta with increased discrimination. **(A)** Specifying a minimum number of pixel overlap of 6 or more, removes some overlapping puncta (arrow), while other puncta remain (arrowhead), in this image of anti-bassoon (magenta) and anti-gephyrin (cyan) from BLA. Scale bar is 10 μm for the panels in A. **(B)** In this image of anti-bassoon from SNc, applying the subtract background with a radius of 10 efficiently removed background staining while not modifying the brightly labeled puncta. Scale bar is 10μm, and applies to panels in B-D. **(C)** Original immunostaining for NR1 (cyan), tyrosine hydroxylase (blue) and YFP in SNc from the same area as the bassoon in panel B. **(D)** The merge images for the original staining (left), identified overlapping anti-bassoon/anti-gephyrin puncta from the entire panel (center), and identified overlapping YFP^+^ PPN axonal anti-bassoon puncta with TH^+^ SNc neuronal anti-NR1 puncta (right) demonstrate the utility of SynapseJ in identifying neuronal sub-class limited expression of synaptic markers.

The macro may optionally correct for decreases in intensity between slices in a Z-stack. These changes in intensity may be due to incomplete penetration of antibodies or bleaching of the signal due to high intensity laser excitation of previous positions in the Z-stack. The user is prompted for the number of slices and can type in a correction factor that realigns the intensities across the sections.

Finally, in some cases the staining of tissue may induce a background hue in the image preventing easy delineation of puncta from the background staining. Using the ImageJ/FIJI processing technique, “Subtract Background”, the user may specify an appropriate radius for a rolling ball filter. The user is capable of applying this to either the pre- or post-synaptic channel with a unique radius for each channel. Immunostaining for bassoon in substantia nigra pars compacta (SNc) had just such a background (Fig. 8B). Using a radius of 10, the filter efficiently removes this background, which maybe non-uniform in intensity, leaving the puncta untouched (Fig. 8B). This process leads to an increase in the signal to noise ratio, allowing for more accurate detection of puncta.

### Delineating synapses localized to labeled neurons

One of the advantages of confocal microscopy is the multiplex labeling of the tissue, allowing for additional qualification of both the pre- and post-synaptic stains to specific cell types. We tested this type of labeling by labeling a synaptic pair of neurons. Mice were bilaterally, stereotaxically injected with an AAV encoding channelrhodopsin tagged with yellow fluorescent protein (YFP) to label pedunculopontine tegmental nuclei (PPN) neurons (Fig. 8C). PPN neurons innervate the SNc dopaminergic neurons. Dopaminergic neurons require expression of tyrosine hydroxylase (TH), the rate-limiting enzyme in synthesis of dopamine that is localized to the cytoplasm (Fig. 8C). Thus, YFP-expression will be used to demarcate Bassoon staining, while immunostaining for TH will be used to label dopaminergic neuron expression of an obligatory subunit for N-methyl-d-aspartate receptors, NR1.

Similar to finding overlapping puncta expression, the macro removes bassoon puncta that do not overlap YFP and removes NR1 expression that do not overlap TH (Fig. 8D). YFP and TH positive neurites and cell bodies are localized by having a minimum intensity. Additionally, we selected puncta as being positive for the marker, by measuring the marker intensities that overlapped with the synaptic puncta. Only puncta having both a minimum intensity above threshold, suggesting the puncta fully overlaps the marker, and a maximum intensity above an intensity, which suggests puncta are well-localized over the most intensely labeled part of the cell, are maintained for further analysis. These puncta are culled before attempting to find puncta that are overlapping between the two synaptic channels.

Using this technique we can reliably identify Bassoon puncta overlapping with YFP positive PPN efferents innervating the SNc. Similarly, we can reliably identify NR1 puncta that are localized to TH-positive, dopaminergic neurons of the SNc. Together, these markers identify putative glutamatergic synapses between PPN glutamatergic neurons and SNc dopaminergic neurons (Fig. 8D). Thus, we can compare all putative glutamatergic synapses in the SNc to the PPN-innervated glutamatergic synapses (Fig. 8D). This novel type of analysis is ideal for detecting changes in neural circuitry due to environmental or genetic manipulations.

## 4 Discussion

We have described a new, open-source, free macro developed on top of the popular ImageJ/FIJI interface to identify overlapping pre- and post-synaptic puncta. The routines that are used by the macro are well-characterized functions of the ImageJ interface, which efficiently identifies puncta in either sparsely- or densely-labeled images. Through a combination of size exclusion and overlap with the alternate channel, we are able to identify a wide array of size and synaptic arrangements, including multi-labeled pre-synaptic puncta. While the identification of synapses is ultimately the goal, the power of the correlation analysis can not be understated. By directly identifying the puncta that are overlapping, detailed morphometric analysis of how the pre- and post-synaptic puncta are correlated or not is a feature distinct to this macro. Image output from the macro allows for demonstration of all collected puncta and the puncta that are overlapping puncta from the alternate channel. Additionally, the lists of ROI can be reentered into the ROIManager to allow for detailed tracing of each puncta from the lists.

The macro requires user input only at the beginning, and once locations for input image files and for output files and parameters are set, the macro proceeds through each image. Progress is denoted image by image and by the final line exclaiming the macro is ready for more. As a brute force method, large images with dense numbers of puncta may very well take a few hours to process per image. The speed of analysis is hastened as the number of raw puncta are excluded for not overlapping puncta from the alternate channel. Increasing clock speed or use of a graphical chipset to handle arithmetic operations would lead to further increases in speed. However, the alternative of hand analysis or use of electron microscopy would not yield the same quantity of data in the time allotted.

One question is how well does SynapseJ do in identifying synapses? One source of validation is in the proportion of inhibitory synapses found in the four regions analyzed in this paper. In adult animals, bassoon is generally localized to presynaptic release sites, such that the number of bassoon puncta would be an approximate measure of all of the synapses (Fig. 2D-H), whereas gephyrin-positive bassoon puncta is a measure of inhibitory synapses (Fig. 4B, C) (Craig et al., 1996;Richter et al., 1999;Micheva et al., 2010;Groeneweg et al., 2018;Pizzarelli et al., 2020). For the BLA, the macro found ~50 bassoon puncta per 100μm^2^, which fell to ~14 gephyrin-positive bassoon puncta per 100μm^2^ (Fig. 2D and 4B). For the VMH, the macro found ~30 bassoon puncta per 100μm^2^, which fell to ~2.5 gephyrin-positive bassoon puncta per 100μm^2^ (Fig. 2D and 4B). For the Arc and CeA, the macro found ~15 bassoon puncta per 100μm^2^, which fell to ~2 gephyrin-positive bassoon puncta per 100μm^2^ (Fig. 2D and 4B). The BLA has a significant inhibitory projection from CeA in addition to a few classes of local inhibitory interneurons, suggesting that an inhibitory percentage at 25-30% of all synapses is approximately correct (Muller et al., 2006;Perumal et al., 2021). While the total number of synapses varies for Arc, CeA, and VMH, each region has an apparent inhibitory synapse percentage of 13-9% of all synapses. All three locations have significant excitatory inputs to the region with a smaller set of inhibitory interneurons or inputs, suggesting these percentages are within reason (Nishizuka and Pfaff, 1989;Andermann and Lowell, 2017;Hashikawa et al., 2017;Babaev et al., 2018;Fadok et al., 2018). Furthermore, gephyrin is not a marker for all inhibitory synapses, so there may be a number of synapses missed by this analysis (Groeneweg et al., 2018;Pizzarelli et al., 2020).

While there is room for error of over and under assignment of putative synapses, we suggest that these errors will be consistent between images. Use of the “Find Maxima” function for sparsely labeled images does not hinder analysis, so if there was a dramatic loss of a synaptic marker, such that what was once a densely labeled image became sparse would not lead to an alteration of the results. This macro would be able to identify whether the remaining puncta were still synaptic or simply residual non-localized expression. Over and under counting of putative synaptic puncta can be altered by adjusting the degree of overlap between puncta of the two channels. There are additional tools available to adjust the included puncta by examining the distance between the centers of the puncta. The center of the puncta is defined as both the geometric center which ignores any heterogeneity in the staining or as the center of mass defined by the distribution of intensity values within the punctum. Either of these distance values identify puncta that are closer and thus more likely to be synaptic.

Of note is the consistent changes that occur in size, intensity and integrated density both as the threshold intensity and minimum sizes were changed, but also as the bassoon or gephyrin puncta are classified by being gephyrin-positive or bassoon-positive, respectively. The direct correlation of changes between mean puncta intensity and threshold intensity, with a reduced but indirect correlation between puncta area and threshold intensity, suggests that a straightforward increase in puncta due to decreasing threshold intensity. However, the direct correlation of changes between puncta area and minimum size, but non-existent correlation between mean puncta intensity and minimum size suggest that the interaction of size and intensity is not simple. Additionally, the average of puncta mean intensity and puncta area are both elevated when comparing the original list of puncta to puncta that are overlapping (Fig. 5A and B, D and E, 6A and B, or D and E). The distribution of puncta sizes and intensities also significantly shifts as puncta are classified based on overlap (Fig. 7A, B). Thus, while the macro robustly identifies puncta of varying sizes and intensities, the overlapping puncta are larger and brighter, which would be expected for a synapse.

Although false positive puncta may be easy to eliminate, reclaiming false negative puncta may be harder. The false negative puncta would be puncta that should be a synapse but were removed anyways. This type of error is harder to address, but one way is to analyze Z-stacks instead of simply a single image plane. The orientation of a pre- and post-synaptic puncta in three dimensions is collapsed into a single imaging plane as two circles that could be completely overlapping if they are oriented perpendicular to the imaging plane or as two discrete circles if they are oriented within the imaging plane. In the perpendicular orientation, as the focal plane is changed to capture the Z-slices, one slice may contain only one of the two synaptic markers. However, as you change your focal plane, a subsequent plane should show complete overlap with a deeper plane now only showing the second synaptic marker. If the puncta are localized within the plane, there may not be enough overlap of the two puncta. However, based upon the diffraction limit of light, use of conventional, non-super-resolution microscopy would lead to overlap to the synaptic markers for both the in-plane orientation and in the perpendicular orientation.

The utility of this macro to output both quantitative data and graphical data is critical. The graphical results with associated lists of ROIs enables visual inspection of the results. This visual inspection is to ensure that puncta were accurately identified and may suggest issues with the parameter settings chosen. Additionally, these images may be further analyzed graphically for non-random distributions of synapses. In many brain regions, the larger inhibitory synapses are located closer to the cell body. One could consider pseudo-coloring ROIs based upon size and examine distance to a nucleus and may find that the puncta closer to the cell body are in fact larger. Further, the identification of puncta that are overlapping cell-specific markers enables the identification and analysis of neural circuits. Ideally, this data would be used to corroborate data garnered from other functional experiments, such as electrophysiology.

An important consideration is how this image analysis procedure compares to other methods. One ImageJ plugin that has some similar features is “Synapse Counter” (Dzyubenko et al., 2016). The pre-processing of the images is remarkably similar, though the user has less choice as to which routines are applied in “Synapse Counter”. The “Synapse Counter” plugin requires very little user input, relying on the range of intensity values within the image to determine and remove background and the threshold intensity. One concern of relying on auto-thresholding and removal of background is this could lead to uneven application of these parameters between images that may vary due to real biological differences. The routine also does not correlate regions of overlap, relying on only counting these regions. During processing, the use of a maximum filter with a 1-2 pixel radius, which aids in identification of puncta, causes significant blurring of the original, obscuring more detailed morphometric analysis. The strength of the SynapseJ routine lies in correlating the observed overlapping puncta and the detailed morphometric output, allowing for more than just counting of the overlap. The extension of SynapseJ to identify synaptic co-localizations with markers of cell types is also unique, and allows for the application to identify and measure synapses at defined points within a neural circuit.

## 5 Author Contributions

GB and JM conceived of and designed the study. JM, PV, KW, SR, and AK performed experiments and collected data. GB, JM, PV, and KW coded the macro. GB wrote the first draft of the manuscript. All authors contributed to manuscript revision, read, and approved the submitted version.

## 7 Acknowledgments

We thank the entire Beaudoin lab for comments and support for this project. Extensive beta-testing of the software package was provided by the Neurobiology classes taught at Trinity University Fall 2016-Fall 2020.

## 8 Software Availability

SynapseJ is available from: https://github.com/GB3Trinity/SynapseJ

